# Multi-Stage Feature Selection (MSFS) Algorithm for UWB-Based Early Breast Cancer Size Prediction

**DOI:** 10.1101/2020.02.06.936831

**Authors:** V. Vijayasarveswari, A.M. Andrew, M. Jusoh, T. Sabapathy, R.A.A. Raof, M.N.M. Yasin, R.B. Ahmad, S. Khatun

## Abstract

Breast cancer is the most common cancer among women and it is one of the main causes of death for women worldwide. To attain an optimum medical treatment for breast cancer, an early breast cancer detection is crucial. This paper proposes a multistage feature selection method that extracts statistically significant features for breast cancer size detection using proposed data normalization techniques. Ultra-wideband (UWB) signals, controlled using microcontroller are transmitted via an antenna from one end of the breast phantom and are received on the other end. These ultra-wideband analogue signals are represented in both time and frequency domain. The preprocessed digital data is passed to the proposed multi-stage feature selection algorithm. This algorithm has four selection stages. It comprises of data normalization methods, feature extraction, data dimensional reduction and feature fusion. The output data is fused together to form the proposed datasets, namely, 8-HybridFeature, 9-HybridFeature and 10-HybridFeature datasets. The classification performance of these datasets is tested using the Support Vector Machine, Probabilistic Neural Network and Naïve Bayes classifiers for breast cancer size classification. The research findings indicate that the 8-HybridFeature dataset performs better in comparison to the other two datasets. For the 8-HybridFeature dataset, the Naïve Bayes classifier (91.98%) outperformed the Support Vector Machine (90.44%) and Probabilistic Neural Network (80.05%) classifiers in terms of classification accuracy. The finalized method is tested and visualized in the MATLAB based 2D and 3D environment.

## Introduction

The rate of a woman contracting breast cancer is reported at a worrying rate globally, particularly in developing countries. Symptoms of breast cancer, for instance, visual changes in the breasts are usually discovered only at the final stage [1]. Consequently, most of the breast cancer cases are detected in the latter stage, at which, are deemed as too late for medical treatment, thus causing death [1–2].

Malaysian National Cancer Registry (NCR) Report published every 5 years recorded that the breast cancer is the most common cancer type, holding the top position out of the other common cancer type [3]. The report also states that the Age-Standardized Incidence Rate (ASR) for female is 34.1 per 100 000 populations in the year 2012-2016. Age-Standardized Incidence Rate for male is also recorded, at the increased rate of 0.5 per 100 000 population [3].

In another report generated by GLOBOCAN in 2018, states that breast cancer holds record as second most commonly diagnosed cancer type in the world, with 2.089 million incidences of reported new cases (11.6%) [4]. Based on the reports, it can be clearly concluded that breast cancer cases are increasing every year and it is still recorded as second top causes of the woman’s death [4,41].

There are many existing clinical methods in diagnosing and detecting breast cancers. Common diagnostic methods are mammography, magnetic resonance imaging (MRI) and ultrasound scans [5, 34–36]. However, these methods are proven costly, bulky, invasive and are unable to detect the early stages of breast cancer. These limitations are the main barriers for an efficient early breast cancer detection. Detection of breast cancer in the early stage is very crucial for further medical diagnostics and treatment. Slow detection is indirectly reducing the survival rate of the patients [5].

Taking into consideration all the limitations of the conventional diagnostic methods, microwave based ultra-wide-band (UWB) imaging technology can be a potential and promising method for early breast cancer detection as it is convenient, non-invasive, secure and low-cost [6,34,37,38]. The research involving UWB imaging for breast cancer started effectively in year 1999 by S.C. Hagness from Winconsin University, USA [7], and since then, started to gain popularity among the researchers, credit to the advancement of computational power since late 1990s. Basically, researchers used either real-time machines such as a vector network analyzer or machine learning to analyze the UWB signals [1].

Talking about the analysis of UWB signals using machine learning methods, the capabilities of the classifier model are dependent on the features fed into the machine learning model for the training purpose. Better is the feature, higher will be the classification rate of the classifier model. In general, the features are selected based on the different feature selection methods proposed by various researchers in breast cancer size detection. Researchers normally identify their features by mean of feature extraction, feature selection or feature dimensionally reduction methods [1]. Previous researchers depict use of conventional feature selection method, basically, by using a single-stage feature selection method [1]. In single-stage feature selection method, the important features are extracted from the raw data, and the extracted data is further filtered to select only important and useful features. The method is having some drawback where there can be misclassification due to the deficiency of quality data during the feature extraction stage. The exploration and exploitation of the data will be insufficient during the feature selection as the features are reduced at the initial stage. As a result, only some redundant features are selected, and some useful features are lost due to poor data management [8–9].

The proposed Multi-Stage Feature Selection (MSFS) can be a solution in overcoming the mentioned drawbacks of single-stage feature selection method. MSFS can increase the learning model performance in breast cancer detection application [8–9]. The proposed multi-stage feature selection method will be discussed comprehensively throughout the paper. The performance of the proposed method is validated using statistical and machine learning approaches.

## Materials and Methods

In this section, the breast cancer sampling technique, the feature extraction from UWB sensors using various data normalization techniques, the proposed feature selection algorithm, and the classification stages are explained.

Fig 1 shows the flowchart of the overall experimental process involved in this research. The process started with data collection using breast phantoms, and signal preprocessing. Then, it will be followed by the proposed MSFS method which comprises of four stages.

**Fig 1.**
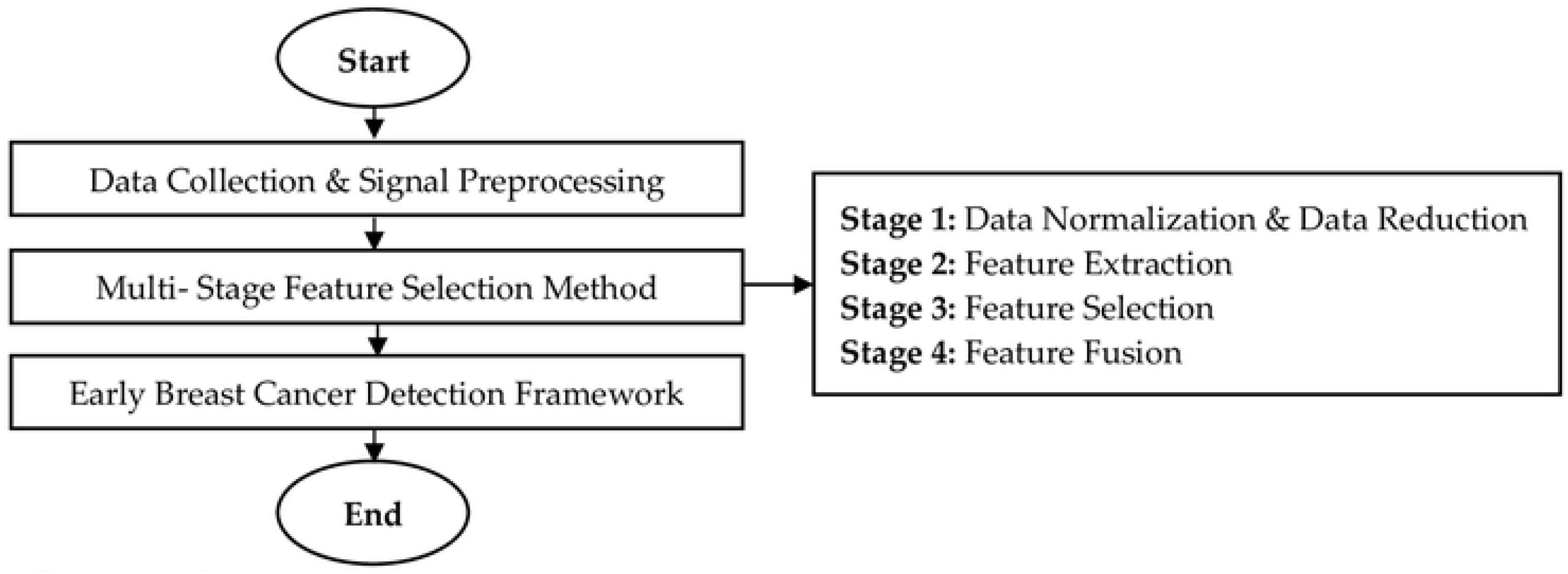
The Overall Experimental Process

In the first stage, the preprocessed data is normalized using various data normalization method, and the data is reduced using Principal Component Analysis (PCA). Data normalization process is important since it is important to select the best features without eliminating useful information from the preprocessed data.

The second stage and all the subsequent stages comprise of feature extraction, feature selection based on the statistical approach, and feature fusion to form the proposed feature that will be incorporated into the Early Breast Cancer Detection (EBCD) Framework. The early breast cancer detection will be visualized in 2D and 3D environment.

### Data Collection

The data collection is conducted using breast phantoms. The breast phantoms have been developed using different materials [10–12]. It is important to make sure that the breast phantoms are having comparable real breast’s dielectric properties in terms of permittivity and conductivity. Based on the literature studies conducted [10–12], most of the researchers use low-cost and non-chemical ingredients like Vaseline (petroleum jelly), a mixture of wheat-flour, water, and soy oil to develop heterogeneous breast phantoms.

The breast phantoms used in this research adopted the same model suggested by the previous researchers [12]. Hemispherical wine glass with 75 mm width, 60 mm height, and 1.9 mm thickness is used as a breast phantom skin. The heterogeneous breast phantom is developed using 100:50:37 ratio of the mixture of petroleum jelly, soy oil, and wheat flour. 25% water is also added to the mixture. Tumors are developed using 10:5.5 mixture ratio of water to the wheat flour. Different tumor sizes are developed for testing (2 mm, 3 mm, 4 mm, 5 mm, and 6 mm). Fig 2 shows the developed breast phantom and tumor for the experiments.

**Fig 2.**
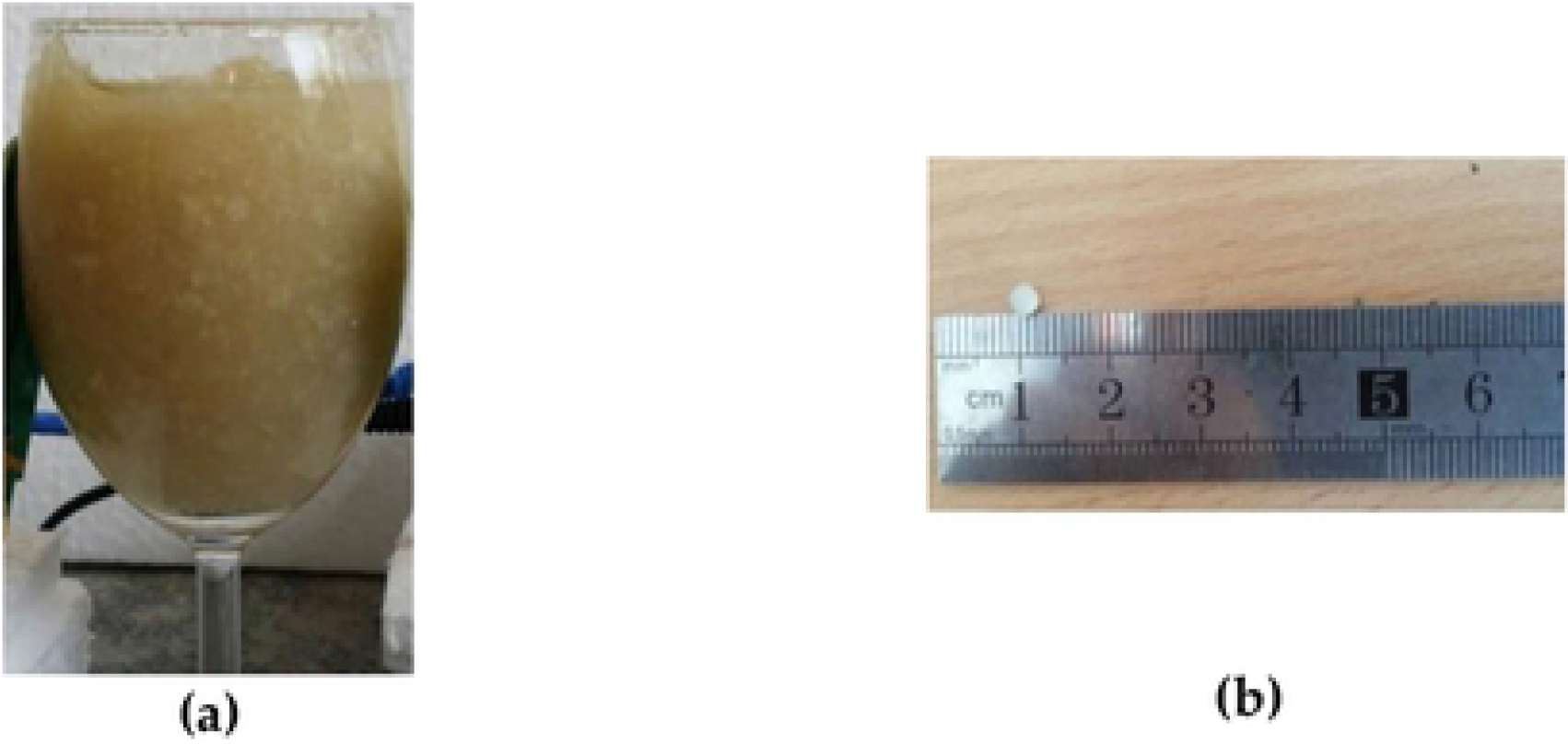
The Developed (a) Breast Phantom (b) Tumor

Fig 3 shows the experimental set-up for the breast cancer sampling [12–16]. A pair of antennae is placed facing each other with the breast phantom located at the middle of the antennae. Feeding cables are used to connect the UWB transceivers with antennae. The UWB signals are generated by the UWB transceivers, passed to transmitter antenna to transmit it on one end and received by the receiver antenna at the diagonal opposite end, concurrently. The captured forward scattered UWB signals are passed to UWB transceiver in the other end.

**Fig 3.**
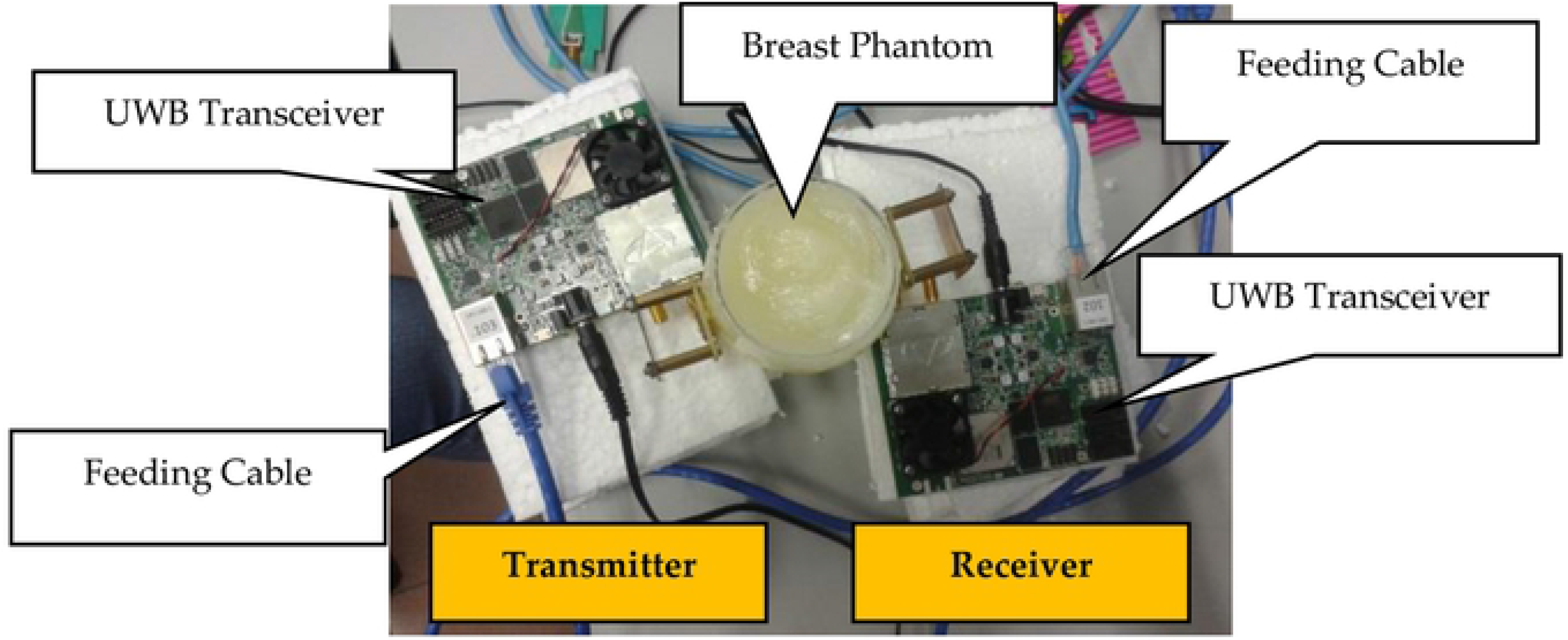
Experimental Set-up for Breast Cancer Sampling

The antennae achieved 6.09 dB gain and 8.15 dBi directivity during the antenna simulation. They are placed close to the breast phantom to avoid any loss of signals and to reduce noises. The UWB transceivers with frequency range of 3 GHz to 10 GHz are used. They are connected to the MATLAB software through Ethernet cross connectors (feeding cables). The receiver antenna captured the forward-scattered signals at the center frequency of 4.3 GHz.

The data collection steps are as follow [12]:

**Step 1:** The 2 mm tumor is placed at starting location in the breast phantom.
**Step 2:** UWB signals are transmitted by antenna and forward scattered UWB signals are captured by the opposite antenna. 50 repetitions are taken at one point.
**Step 3:** The tumor is placed at 27 different locations within the breast phantom. Each tumor (of same size) is placed at different location using the combination location of *x* coordinate (0.25 cm, 2 cm, 3.25 cm, 5 cm and 6.25 cm), *y* coordinate (0.25 cm, 2 cm, 3.25 cm, 5 cm and 6.25 cm) and *z* coordinate (3 cm, 4 cm, 5 cm).
**Step 4: Step 1** to **Step 3** are repeated until all the locations in the breast phantom are covered. The tumor size is then changed. **Step 1** to **Step 4** are repeated until the UWB signals are captured for all five different tumor sizes.

A total of 6750 UWB signals are collected. Each signal sample has 1632 data points. A sample of forward scattered time domain signals (transmitted and received) are shown in Fig 4.

**Fig 4.**
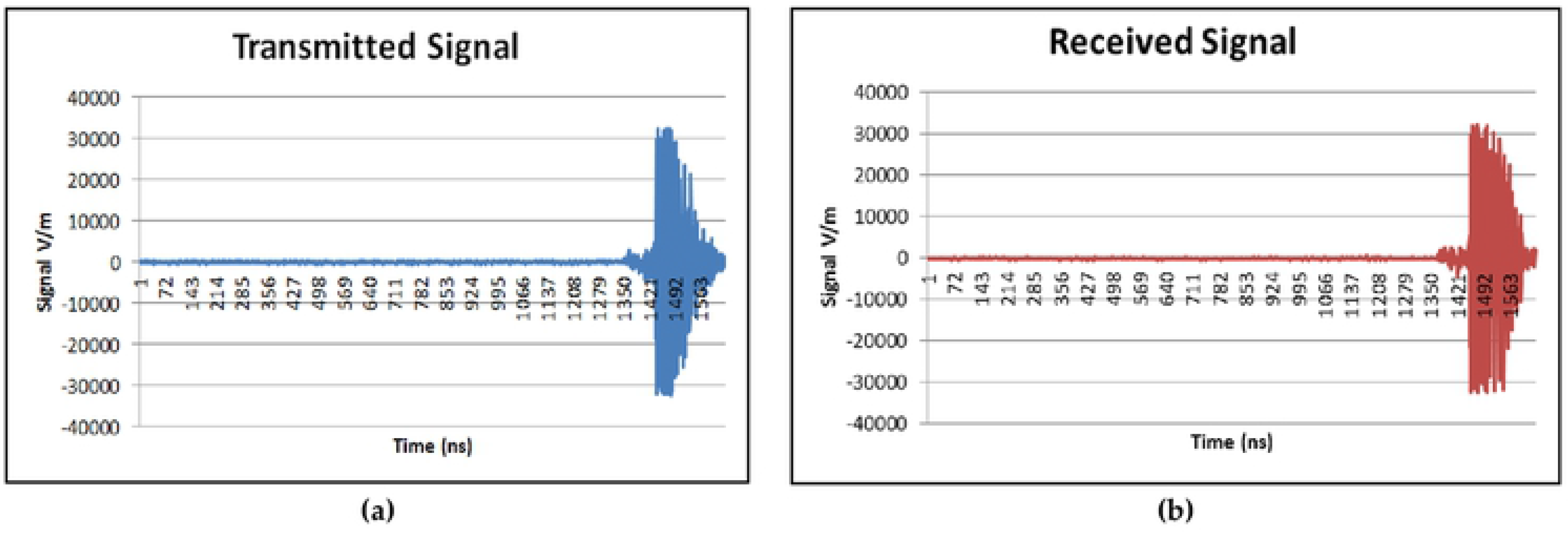
(a) Transmitted UWB Signal and (b) Received UWB Signal

In general, the signal exists in time domain. It is easier to visualize the signal characteristics in time <u>d</u>o<u>ma</u>in. However, analyzing the signal characterization in frequency domain is equally important because it helps to observe the characteristics of the signal which are unable to be visualized in the time domain [17–18,39]. Thus, the time domain signals obtained from the UWB transceivers are transformed to the frequency domain signals using the commonly used Fast Fourier Transform (FFT). Fig 5 demonstrates the representation of the signal in the frequency domain after the transformation. The highest peak of the signal is approximately at 4.3 GHz same as the center frequency of the UWB antenna used.

**Fig 5.**
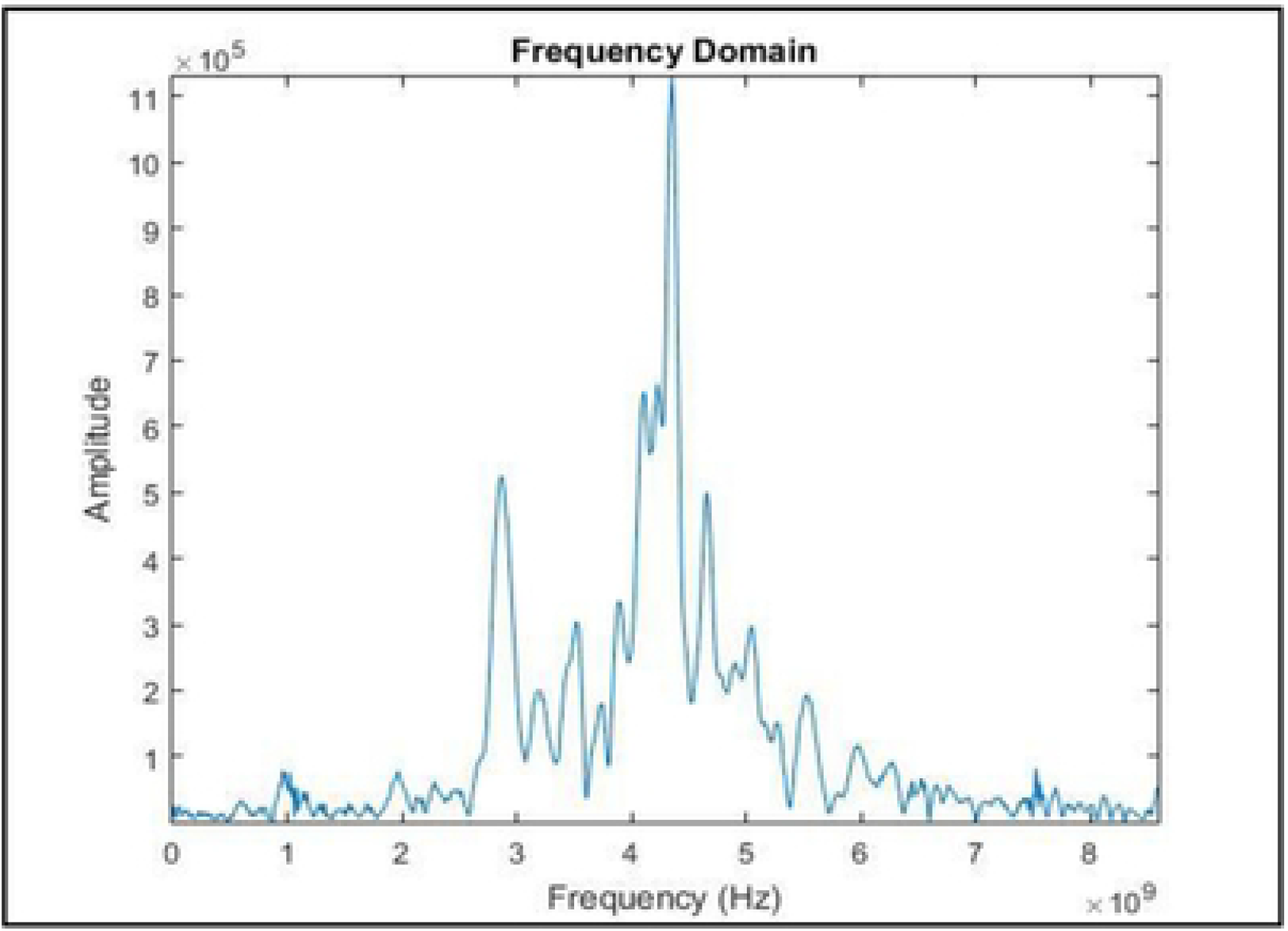
The UWB Signal in Frequency Domain

### Multi-Stage Feature Selection Method

Fig 6 illustrates the overall flow chart of the proposed MSFS method. It is divided into multiple stages [8]. Once the data is preprocessed, it is normalized to 10 different data normalization methods, and the data is reduced using PCA. Then, 10 different features will be extracted from each data normalization method. The best features are selected statistically from the extracted features based on the *p*-value and *F*-value. Then, the selected feature datasets are fused together to produce a newly proposed hybrid feature dataset which will be used for EBCD framework.

**Fig 6.**
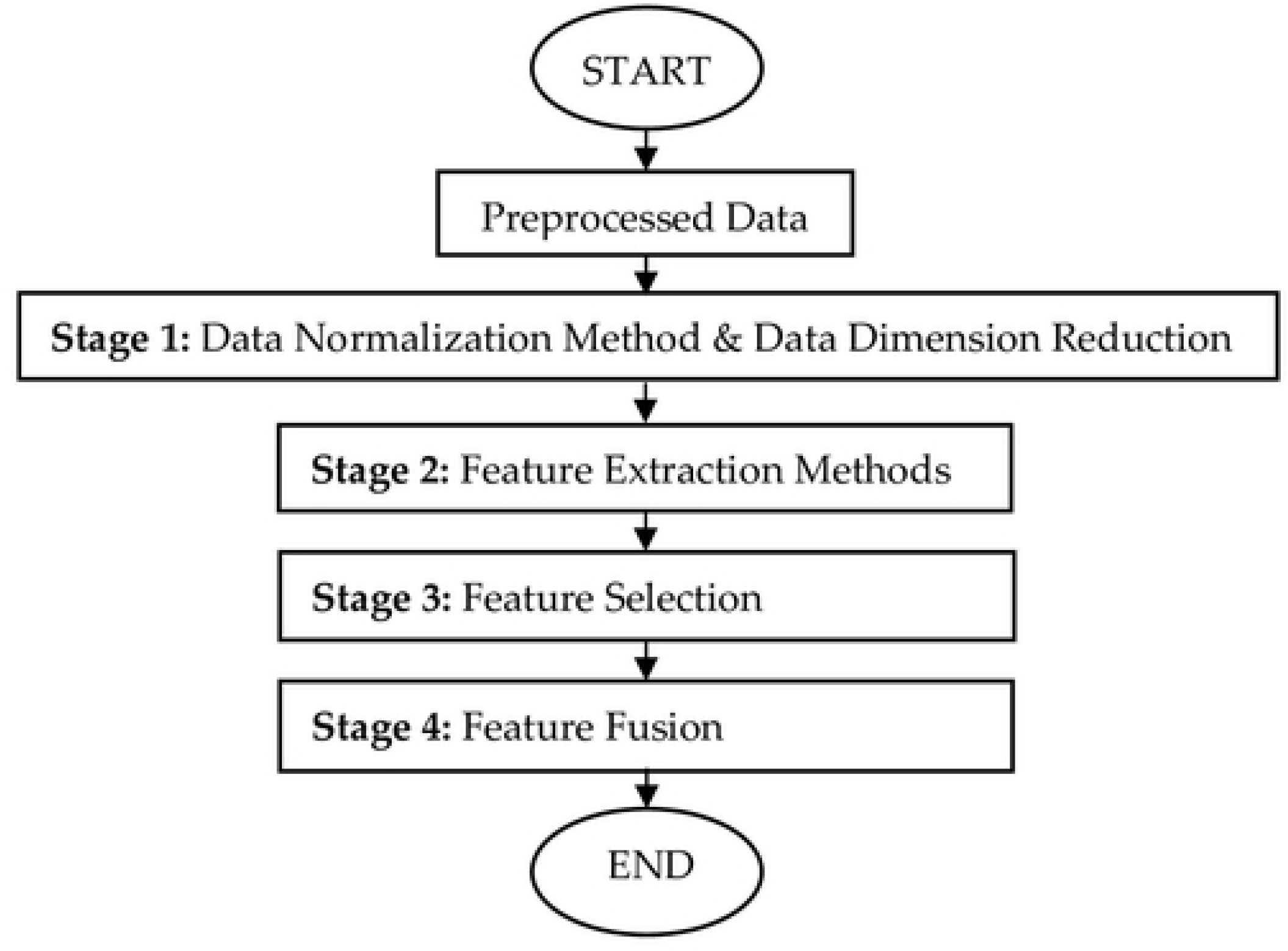
Multi-Stage Feature Selection Method

### Stage 1: Data Normalization Methods & Data Dimension Reduction

Data normalization is a method to standardize the range of features without reducing the dimension of the data [19–22,41]. Data normalization process is important since it is important to select the best features without eliminating useful information from the preprocessed data [19–22]. Conventional single stage feature selection having the drawback of possibly selecting data after eliminating useful data during feature extraction stage. Thus, for this work, raw data samples are normalized using ten different data normalization methods. Based on the comprehensive review done on the previous researches, five data normalization methods are chosen from the commonly used methods, namely, Decimal Scaling (DS), Z-score (ZS), Linear Scaling (LS), Min-Max (MM) and Mean & Standard Deviation (MSD) methods [19–22]. The other five data normalization methods are newly introduced in early breast cancer detection application, namely, Relative Logarithmic Sum Squared Voltage (RLSSV), Relative Logarithmic Voltage (RLV), Relative Voltage (RV), Fractional Voltage Change (FVC) and Relative Sum Squared Voltage (RSSV) [8–9]. These data normalization methods are proposed by [8–9] to overcome the baseline drift error that normally comes together with the data sample which affects the quality of the data samples. The received signals are in amplitude (V/m) versus time (ns) for time domain (refer Fig 4) and amplitude (V/m) vs frequency (Hz) for frequency domain (refer Fig 5). The amplitude (V/m) value is used as voltage input for these five data normalization methods.

Once the data is normalized, the normalized data is dimensionally reduced to remove redundant and statistically insignificant data [21]. The dimension of data is reduced as follows:

a. Each dataset has the maximum number of columns, *c* of principal components.
b. The last column of the principal component is reduced from the dataset. The remaining column is *c*-1.
c. The *p*-value is computed for the *c*-1 column of the principal components.
d. Step 2 and 3 are repeated until the *p*-value is less than 0.05.
e. The remaining number column of the principal components is recorded.
f. These processes are repeated for all ten new normalized datasets.

### Stage 2: Feature Extraction Methods

In order to perform the feature selection process in Stage 3 effectively, feature extraction method is applied on the 10 normalized datasets mentioned in the previous section. Ten features consist of combinations of statistical, time domain and frequency domain features are extracted from each normalized dataset. The features are Mean (M), Standard Deviation (SD), Skewness (S), Variance (V), Power Spectral Density (PSD), FFT Maximum Value (MAX), FFT Minimum Value (MIN), Independent Component Analysis (ICA), Shannon Entropy (SE) and Sure Entropy (SU) [18, 24].

Mean is the ratio of the sum of values to the total number of values as shown in (1).

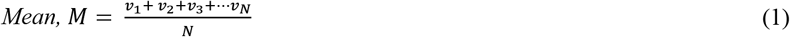

where, *v*_1_ is the first value of data and *N* is the data sample size.

Standard Deviation is used to measure the amount of variation of a set of values in a data as shown in (2).

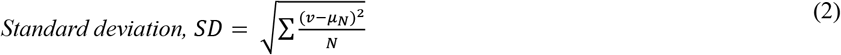

where, *v* is the value of data, *N* is data sample size and *μ_N_* is the mean.

Skewness measures the asymmetry of distribution. Distribution is symmetry if it is looked same for both sides as (3)

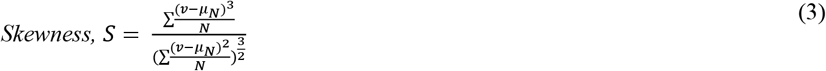

where, *v* is the value of data, *N* is the data sample size and *μ_N_* is the mean.

Variance measures how far the value is from the mean. It is measured using the (4).

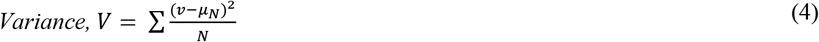

where, *v* is the value of data, *N* is the data sample size and *μ_N_* is the mean.

Power spectral density (PSD) estimates the power in a different frequency range. The time domain data should be transformed into frequency domain data before further analysis. In this study, PSD is estimated using the Welch method which is defined as the moving window technique. Initially, FFT values are computed for each window and then, PSD values are calculated by averaging FFT values over all windows. The Hamming window function is used here because it has a good frequency resolution and reduces spectral leakage [24].

Maximum FFT is the largest value in a set of data after the transformation of time domain data to frequency domain data using FFT. It is usually calculated using the *max* function in MATLAB. It is measured using the (5).

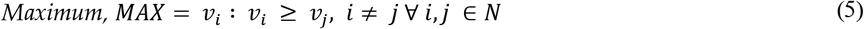

where, *v_i_* is the first value of data, *v_j_* is the second value of data, *i* is 1,2,3 … *i_n_*, *j* is 1,2,3 … *j_n_* and *N* is the data sample.

Minimum FFT is the smallest value in a set of frequency domain data and is calculated using the *min* function in MATLAB. It is measured using the (6).

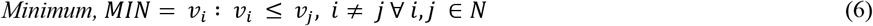

where, *v_i_* is the first value of data, *v_j_* is the second value of data, *i* is 1,2,3 … *i_n_*, *j* is 1,2,3 … *j_n_* and *N* is the data sample.

ICA identifies statistically independent values in a dataset. The equation (7) shows the statistical ICA model.

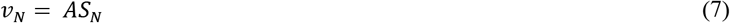

where, *v_N_* is a set of observation vector, *S_N_* is a mixture of independent component vector, *N* is the sample size and *A* is *N* * *N* mixing mixture. Then, ICA finds the unmixing matrix *W* (inverse of *A*) to obtain the independent components (IC) as shown in (8).

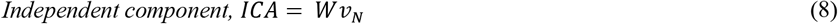

where, *v_N_* is the observation vector and *N* is the data sample size.

Entropy measures the uncertainty distribution and complexity characteristics in data. Shannon entropy (SE) is defined as shown in (9) which describe the internal characteristic information in a data.

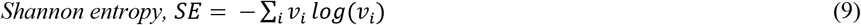

where, *v_i_* is the value of data.

Sure entropy is the measurement of surface entropy and is defined as shown in (10).

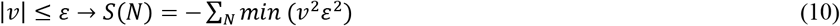

where, *S*(*N*) is sure entropy, *v* is the value of data, *N* is the data sample size and *ε* is the positive threshold which is determined using Steins unbiased risk estimate principle.

### Stage 3: Feature Selection

Stage 3 is divided into two analyses. The first analysis is on selection of normalization method. The second analysis is on selection of features. Both analyses are conducted using statistical computations of statistical *p*-value and *F*-value [8–9].

A. Analysis 1: Selection of Normalization Methods 10 features are extracted from each data normalization method, which resulted total of 100 extracted features. Each normalized method has a data matrix of [6750 x 10] where, 6750 is the number of data samples and 10 is the number of features as in (11) to (20). Each data matrix will be referred with name [DS], [ZS], [SD], [MM], [MSD], [RLSSV], [RLV], [RV], [FVC] and [RSSV] respectively.

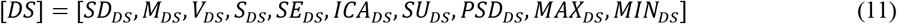

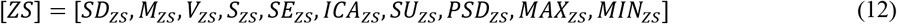

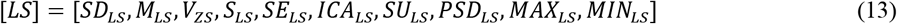

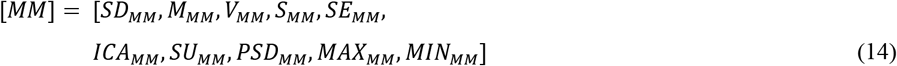

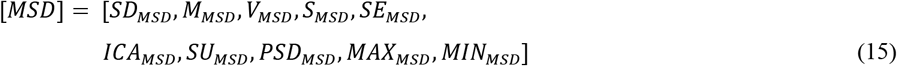

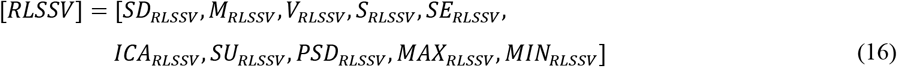

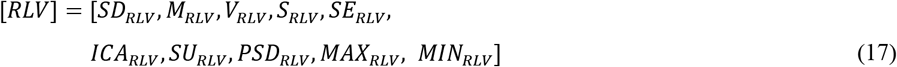

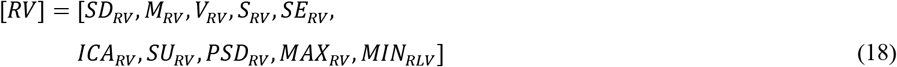

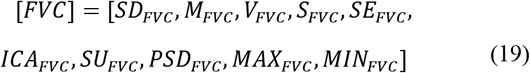

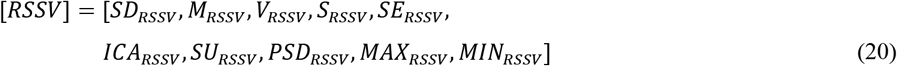 Statistical tests are conducted on each data matrix in (11) to (20) to find the *p*-value and the *F*-value of the respective data matrix. Table 1 shows the *p*-value and *F*-value of the ten data matrix. The five data matrix which pass the selection criteria of having *p*-value less than 0.05 and highest *F*-value are selected. Based on Table 1, it can be seen that data matrices [LS], [MM] and [RLV] did not meet the first selection criterion of (*p* < 0.05), and thus, rejected. The second selection criterion (highest *F*-value) is checked for the remaining seven data matrices, and it is found that [RLSSV], [DS], [ZS], [RV] and [FVC] are selected based on the two mentioned selection criteria. These five data matrices are used for further analysis.
B. Analysis 2: Selection of Feature Extraction Methods The features mentioned in (1) to (10) are extracted using the finalized data normalization methods in Analysis 1. The 50 extracted features undergo the same selection criteria of *p-*value is less than 0.05 and highest *F*-value. If the feature is statistically significant (*p*-value is less than 0.05), then features are selected for second criterion check. Among the statistically significant features, ten best features are selected based on highest *F*-value. Table 2 shows the result of the analysis. The statistically insignificant features (*p*-value > 0.05) are represented with symbol (-), and therefore, are removed from the feature selection listing. Out of the remaining features, ten features with highest *F*-values (highlighted in the table) are selected for Stage 4. Table 3 shows the ranking of the selected features in Stage 3 based on the *F*-values (in descending order) for the respective normalization methods.

**Table 1.**
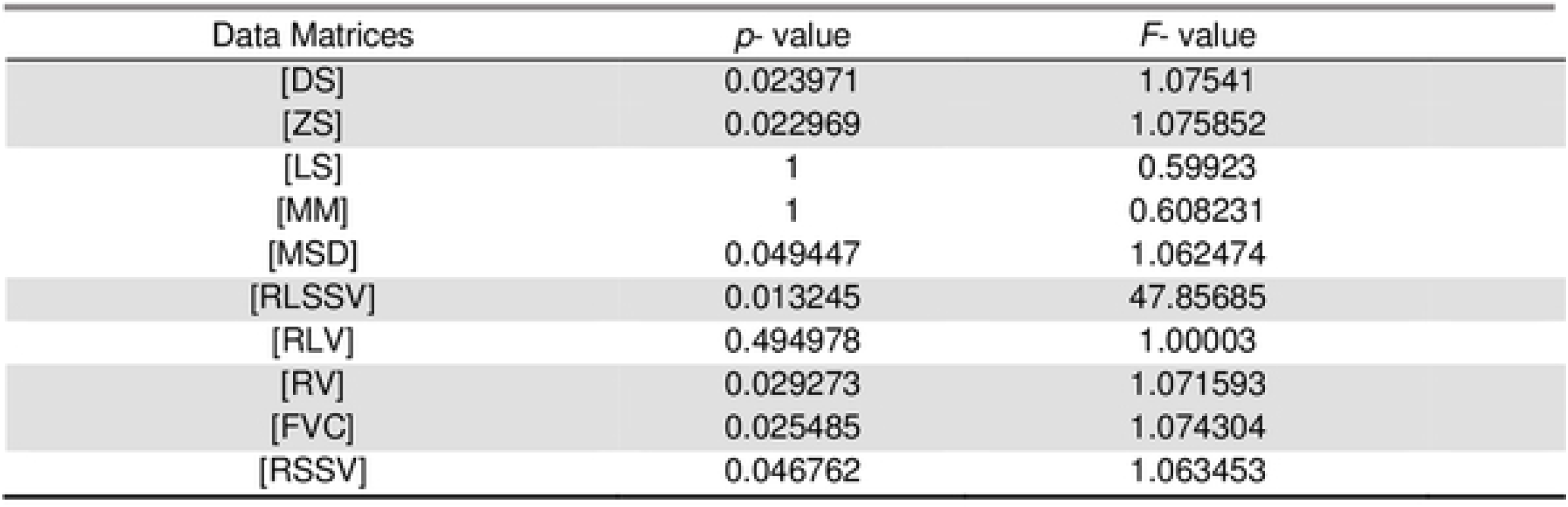
*p*-value and *F*-value Computation for the Data Matrices

**Table 2.** *F*-value of Extracted Features for Finalized Normalization Method

|  |  | Data Normalization Methods |  |  |  |  |
| --- | --- | --- | --- | --- | --- | --- |
|  |  | ZS | DS | RLSSV | RV | FVC |
| Feature Extraction Method | SD | - | 55.12 | 10.95 | 103.64 | 160.47 |
|  | M | - | - | 5.46 | - | - |
|  | V | - | 53.79 | 13.76 | 88.70 | 135.12 |
|  | S | - | - | 126.54 | - | - |
|  | SE | - | 70.59 | 9.48 | 154.30 | 140.44 |
|  | ICA | 8.48 | 2.36 | - | 10.22 | - |
|  | SU | 282.76 | 53.89 | 3.47 | 88.70 | 48.21 |
|  | PSD | 222.52 | - | - | 11.59 | 3.29 |
|  | MAX | 6.76 | 4.26 | - | - | - |
|  | MIN | 4.45 | 37.55 | - | - | - |

### Stage 4: Feature Fusion

Feature fusion is the hybridization of the statistically selected features. Table 3 shows the ranking of the selected features in Stage 3 based on the *F*-values (in descending order). They are used in the further analyses using the Dataset Name assigned.

**Table 3.** Ranking of Selected Features in Stage 3 based on the *F*-value

| Feature | Data Normalization Method | $F$- value | Dataset Name |
| --- | --- | --- | --- |
| SU | ZS | 282.76 | [SU-ZS] |
| PSD | ZS | 222.52 | [PSD-ZS] |
| SD | FVC | 160.47 | [SD-FVC] |
| SE | RV | 154.30 | [SE-RV] |
| SE | FVC | 140.44 | [SE-FVC] |
| V | FVC | 135.12 | [V-FVC] |
| S | RLSSV | 126.55 | [S-RLSSV] |
| SD | RV | 103.64 | [SD-RV] |
| V | RV | 88.70 | [V-RV] |
| SU | RV | 88.70 | [SU-RV] |

In this stage, the selected features are fused together to develop the proposed hybrid feature. Each dataset is reduced to single column [6750 rows x 1 column] using Singular Value Decomposition (SVD) method [8]. Before data fusion, each dataset is having [6750 x 1] dimension as shown in Fig 7. The ten individual datasets are fused together to form the proposed hybrid feature with dimension [6750 x 10]. First column to the last column are assigned based on the ranking in Table 3, starting from [SU-ZS] to [SU-RV].

**Fig 7.**
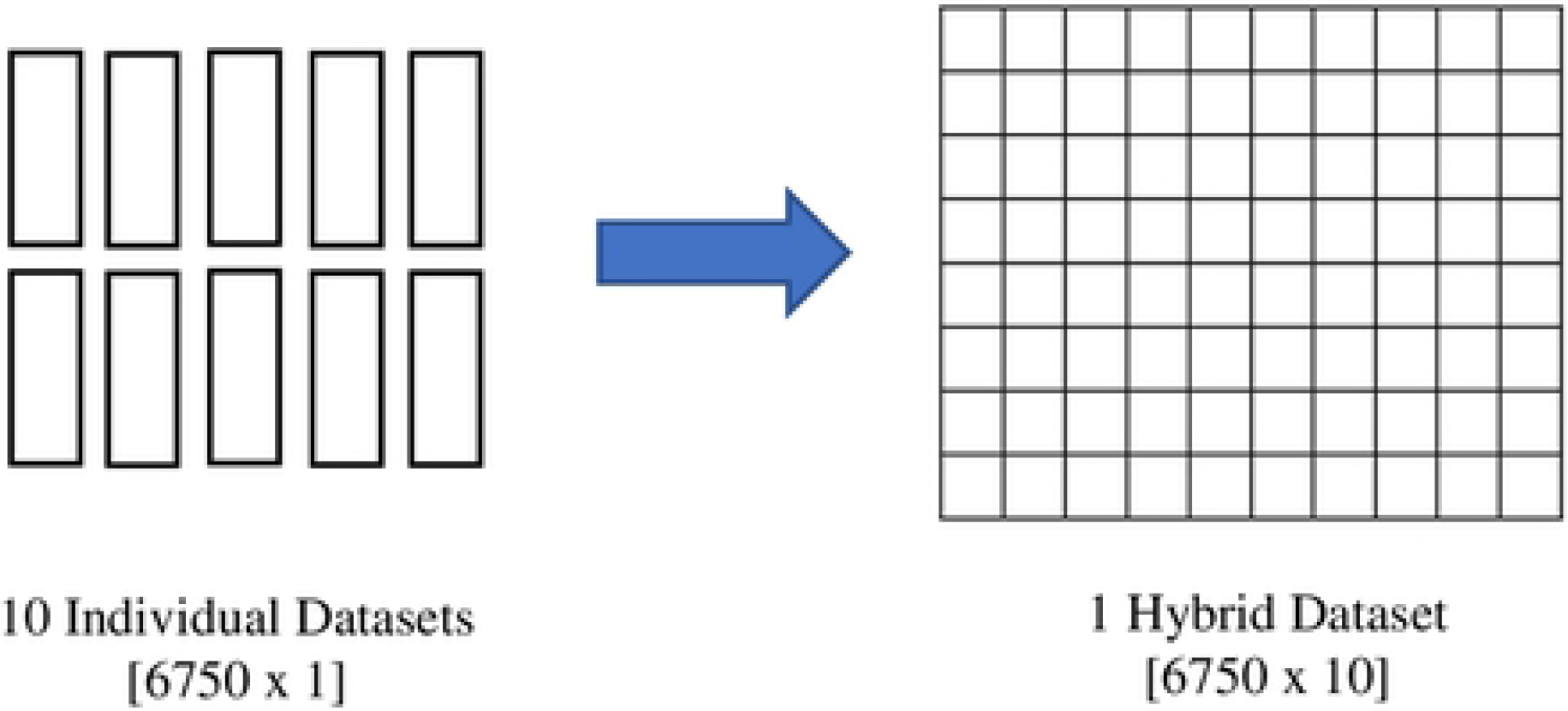
Fusion Process of the Hybrid Feature

*F*-value is computed for different number of hybrid features, starting from fusion of 10 best features, decreasing until fusion of 2 best features. The results are tabulated in Table 4. From the result, it can be seen that fusion of 6 to 10 best features are giving best result range in terms of *F*-value. Fusion of 8 best features recorded the highest *F*-value. Three datasets with highest *F*-value are chosen for further analysis. The three datasets are referred to as 10-HybridFeature, 9-HybridFeature and 8-HybridFeature datasets throughout the paper.

**Table 4.** *F*-value of the Datasets with Different Number of Features

| Number of Hybrid Features | Dimension [Rows x Columns] | *F*- value | Dataset Name |
| --- | --- | --- | --- |
| 10 | [135 x 10] | 938267.28 | 10-HybridFeature |
| 9 | [135 x 9] | 939582.08 | 9-HybridFeature |
| 8 | [135 x 8] | 942111.32 | 8-HybridFeature |
| 7 | [135 x 7] | 935645.64 | - |
| 6 | [135 x 6] | 937341.03 | - |
| 5 | [135 x 5] | 229.30 | - |
| 4 | [135 x 4] | 208.70 | - |
| 3 | [135 x 3] | 168.38 | - |
| 2 | [135 x 2] | 702.17 | - |

The overall block diagram of the proposed MSFS method is shown in Fig 8 and Fig 9. Fig 8 shows the data normalization selection and data dimension reduction, while Fig 9 shows the feature extraction selection, feature fusion and formulation of the proposed feature datasets. The dimension (Rows x Columns) of the datasets is shown in square brackets ([]).

**Fig 8.**
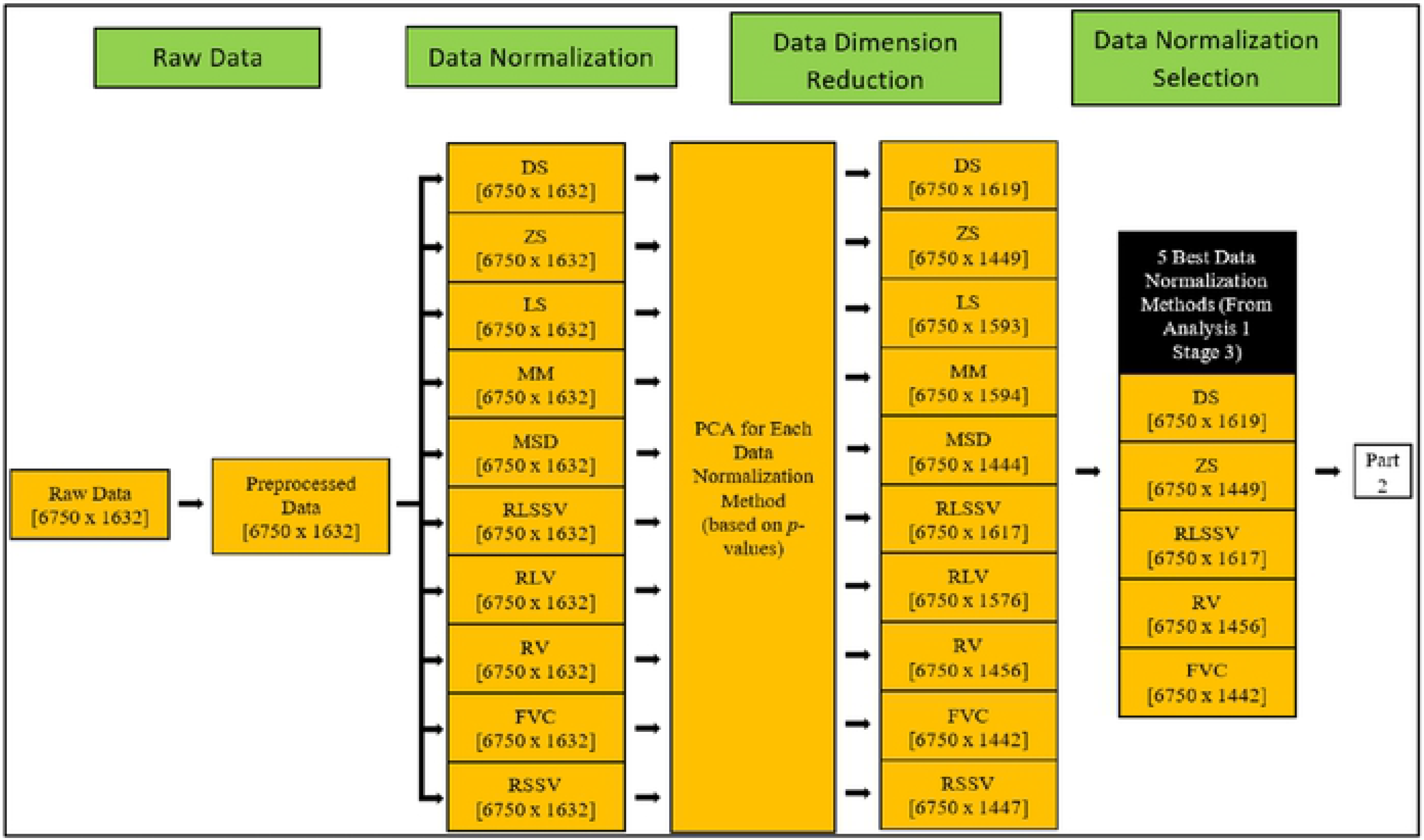
Block Diagram of Proposed MSFS Method (Part 1)

**Fig 8.**
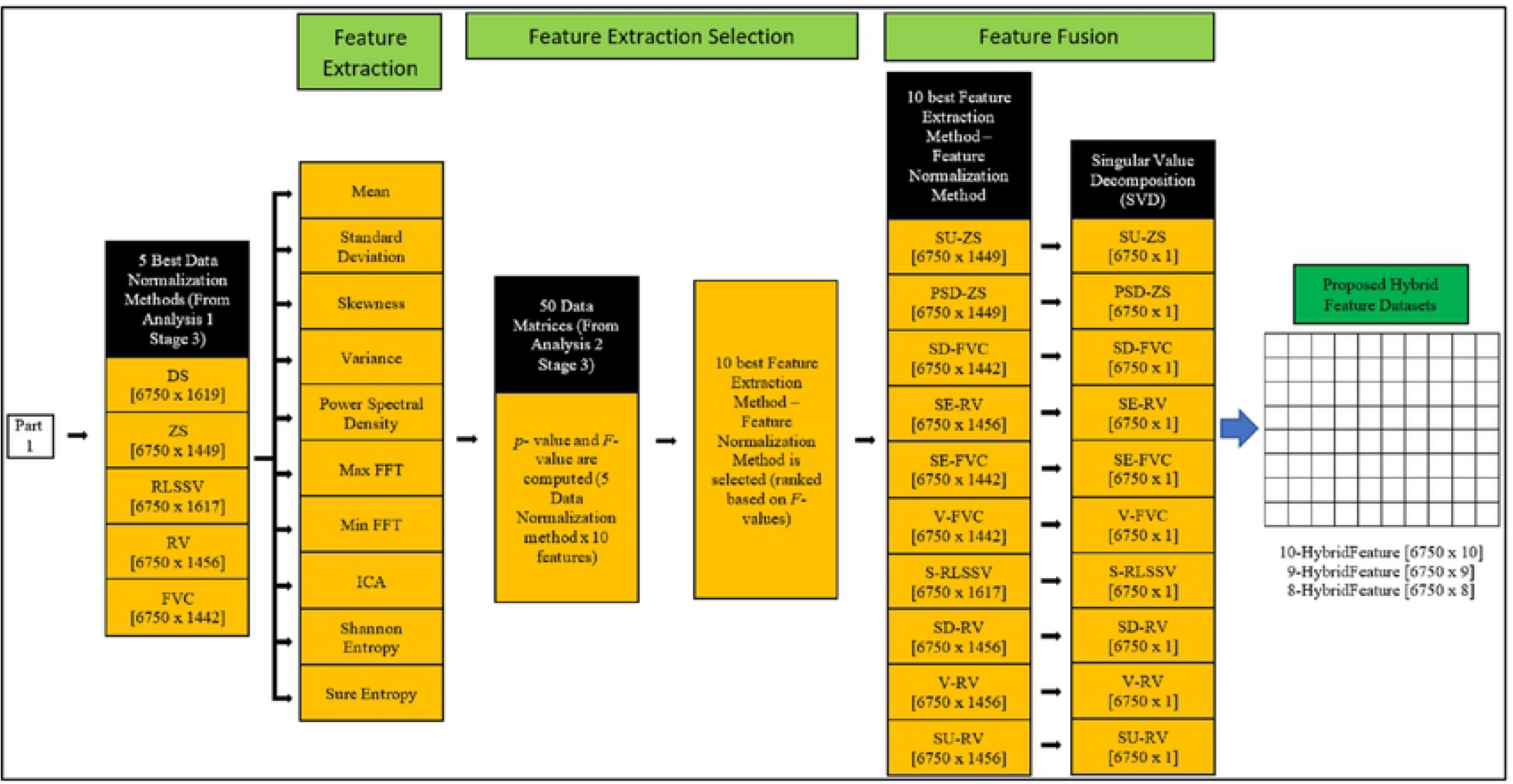
Block Diagram of Proposed MSFS Method (Part 2)

### Classification of Breast Cancer Size

For breast cancer size classification tests with classifiers, the three hybrid feature datasets are used. The SU-ZS dataset from Stage 4 which recorded the highest *F*-value as an individual feature extraction method-feature normalization method is added as a comparison dataset. Three commonly used machine learning methods, namely, Support Vector Machine (SVM), Probabilistic Neural Network (PNN) and Naïve Bayes (NB) classifiers are used for breast cancer size classification [25–31].

The classifier parameters are set in such a way that, for SVM, the linear kernel function is used. For PNN, spread factor of 0.1 and four layers (input, pattern, summation and output layers) are used. There is no classifier parameter for NB since it does not require tuning parameters [29]. There are six possible outputs, which are the five breast cancer tumor sizes (2 mm, 3 mm, 4 mm, 5 mm, and 6 mm) and non-existence of the tumor. Two processes are involved in the classification: training and testing. The training and testing are conducted using k-fold cross-validation method [32]. Fig 10 shows the k-fold cross-validation method used in this research [32–33]. Ten k-folds are used. The total number of 6750 data samples from each dataset are divided into 10 equal portions (folds). Each fold will have 675 data samples. The training process is done using the 9 folds data, while the testing is done using the remaining a fold data. Each fold will take turn to be the testing fold, until the training-testing process completed. Confusion matrices are generated for each iteration, and the accuracy, sensitivity and specificity are calculated for each iteration using equations (21) to (23). The average classification accuracy, sensitivity and specificity of all folds are considered as the performance of the classifier [8].

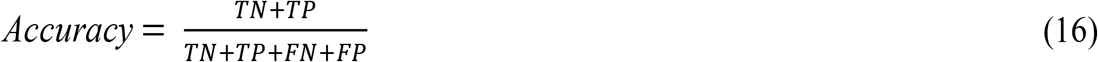

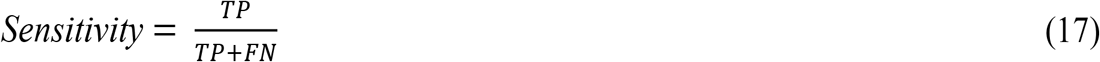

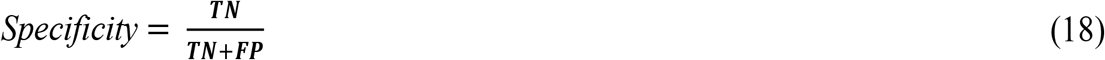

where, *TN* in true negative, *TP* is true positive, *FN* is false negative and *FP* is false positive.

**Fig 10.**
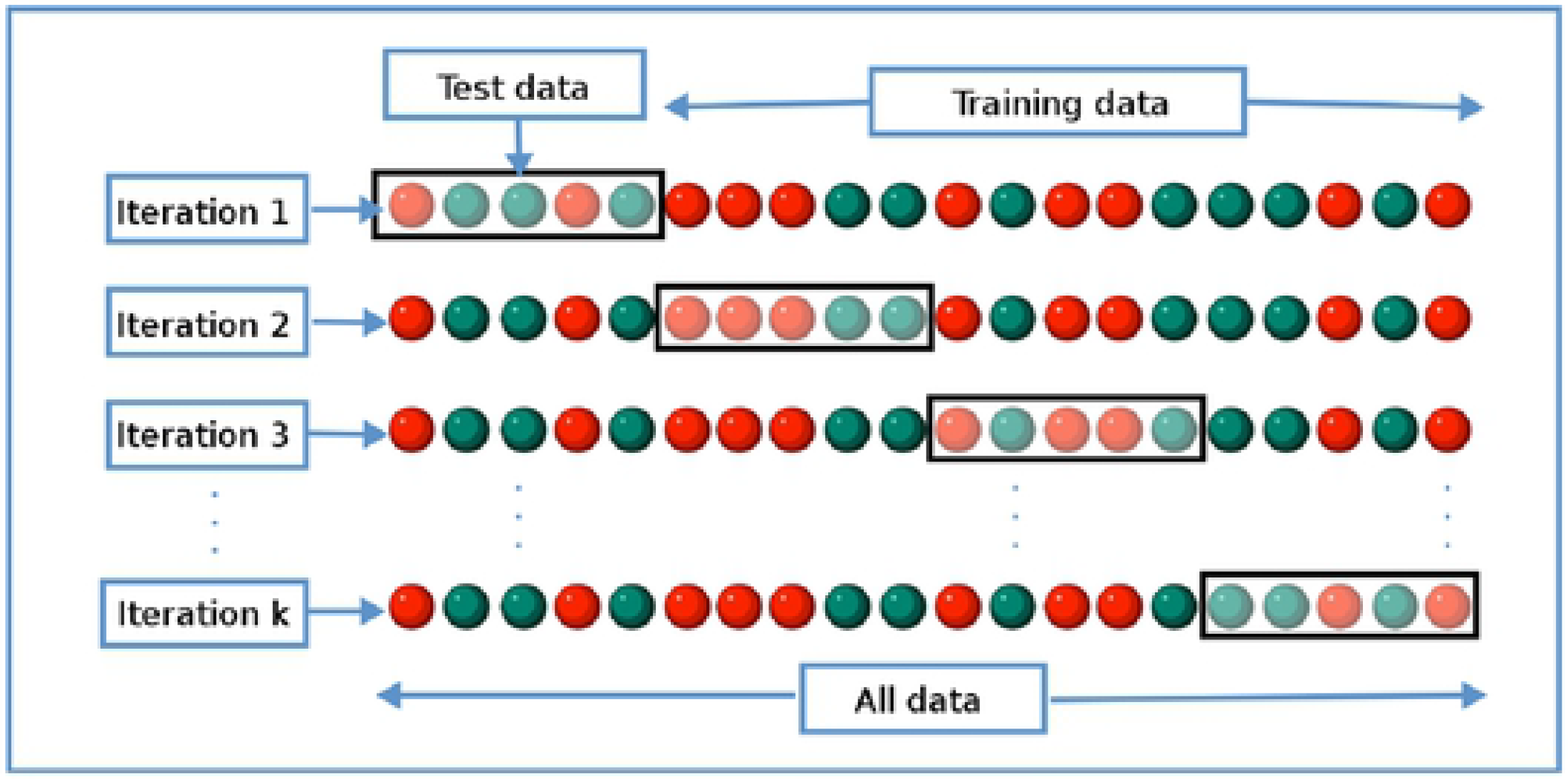
k-Fold Cross-validation Method

## Results

Fig 11 shows the performance of SVM, NB and PNN for SU-ZS, 8-HybridFeature, 9-HybridFeature and 10-HybridFeature datasets. For SU-ZS dataset, the accuracies for SVM, NB and PNN are recorded as 84.39%, 83.69% and 82.97% respectively. These accuracies are considered as the benchmarking accuracies of this study to show the effectiveness of the proposed method. Table 5 shows the performance of SVM, NB and PNN for SU-ZS Dataset (Reference Dataset). 8-HybridFeature dataset records the highest classification performance, whereas, 10-HybridFeature dataset has the lowest classification performance of all classifiers.

**Fig 11.**
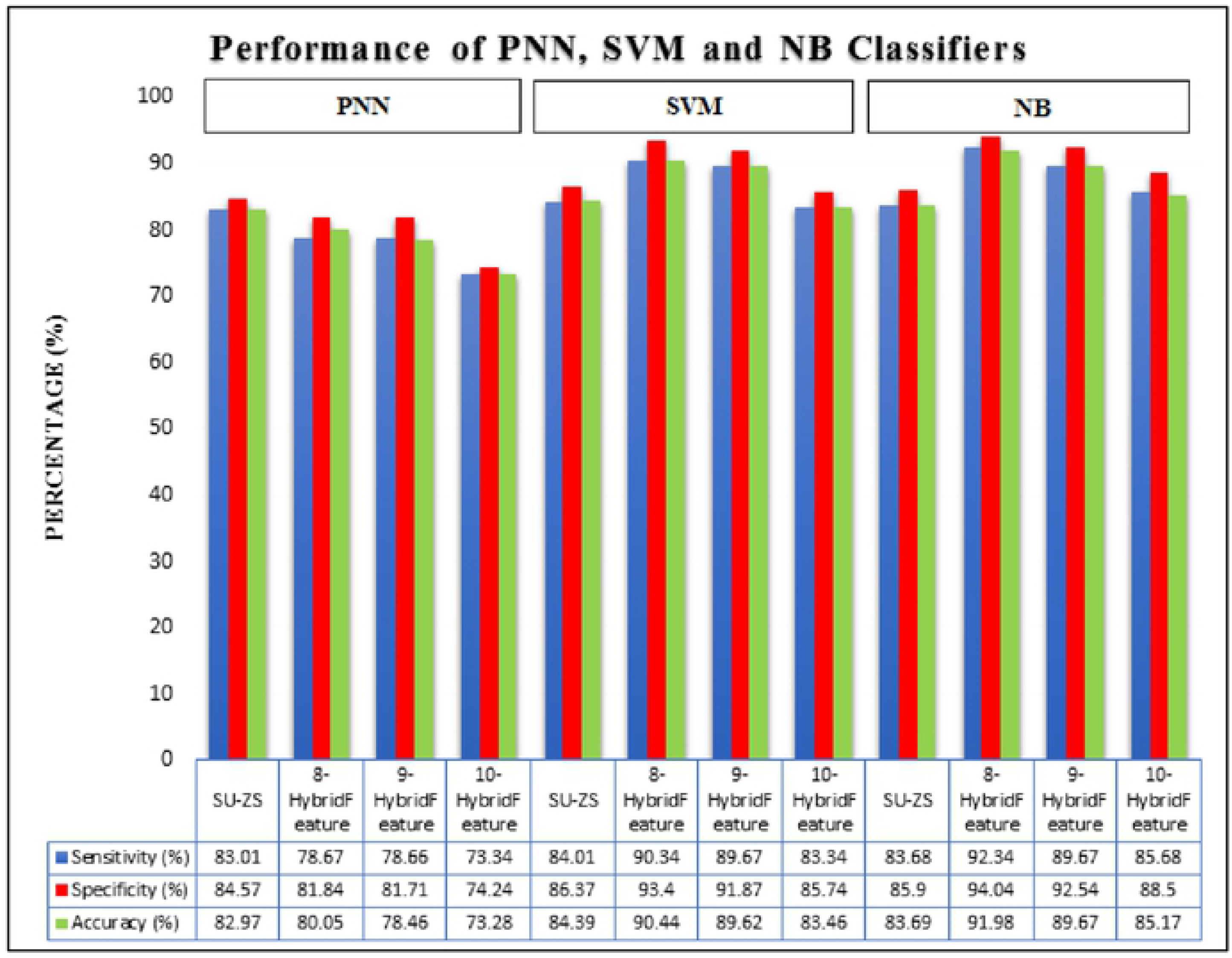
Performance of SVM, NB and PNN for SU-ZS, 8-HybridFeature, 9-HybridFeature and 10-HybridFeature Datasets

**Table 5.**
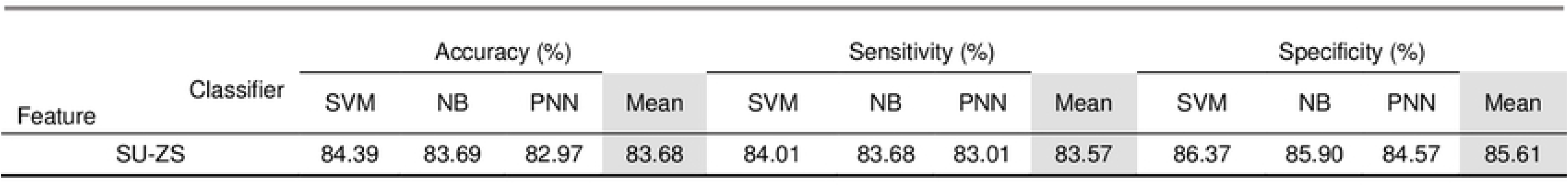
Performance of SVM. NB and PNN for SU-ZS Dataset (Reference Dataset)

| Feature | Classifier | Accuracy (%) |  |  |  | Sensitivity (%) |  |  |  | Specificity (%) |  |  |  |
| --- | --- | --- | --- | --- | --- | --- | --- | --- | --- | --- | --- | --- | --- |
|  |  | SVM | NB | PNN | Mean | SVM | NB | PNN | Mean | SVM | NB | PNN | Mean |
| SU-ZS |  | 84.39 | 83.69 | 82.97 | 83.68 | 84.01 | 83.68 | 83.01 | 83.57 | 86.37 | 85.90 | 84.57 | 85.61 |

For PNN classifier, the highest result is achieved by 8-HybridFeature dataset by obtaining 80.05%, 78.67%, 81.84% for accuracy, sensitivity and specificity respectively. The lowest result is achieved by 10-HybridFeature dataset by obtaining 73.28% accuracy, 73.34% sensitivity and 74.24% specificity.

For SVM classifier, 8-HybridFeature dataset obtains the highest classification performance, which is 90.44%, 90.34% and 93.40% for accuracy, sensitivity and specificity respectively, while the 10-HybridFeature dataset obtains the lowest classification performance with recorded performance 83.46%, 83.34% and 85.74% for accuracy, sensitivity and specificity respectively.

For NB classifier, 8-HybridFeature dataset achieves 91.98%, 92.34% and 94.04 for accuracy, sensitivity and specificity (the highest result). The 10-HybridFeature dataset achieves 85.17%, 85.68% and 88.50% for accuracy, sensitivity and specificity respectively. In general, the classifiers are able to classify with lowest misclassification rate. The three hybrid feature datasets also recorded a better result in general compared to individual features (Table 5) except for PNN classifier. It can be due to unoptimized spread factor for PNN classifier.

8-HybridFeature dataset is proven to be having the best result compared to other datasets because it contains very strong and significant features. It is proven by validation using statistical approach (*p*-value and *F*-value) and machine learning approach. 8-HybridFeature dataset has improved the classification process to be more than 90% accurate for SVM and NB classifiers. Therefore, it can be concluded that the selected hybrid features through MSFS process are able to improve the overall classifier performance.

The accuracy of the proposed method is compared with the existing work, [1,31] as demonstrated in Table 6. The result proves the proposed MSFS method and hybrid feature are better compared to the other existing method. The proposed MSFS method achieves 91.98% which is much better compared to the previous method (74.61%). It improves approximately 17% of accuracy in comparison to the of the previous study.

**Table 6.** Comparison with Previous Studies

| Researcher | [1,31] | Proposed Work |
| --- | --- | --- |
| Method | Extract statistical features | MSFS method |
| Features | 4 (mean, standard deviation, maximum and minimum values) | 8 (8-HybridFeature) |
| Testing Method | 70 % for training, 15% for validation and 15% for testing | k-fold cross- validation (10 folds) |
| Classifier | Feed forward backpropagation neural network | Naïve Bayes |
| Accuracy (%) | 74.61 | 91.98 |

### Early Breast Cancer Detection (EBCD) Framework

The EBCD framework is a user-friendly platform developed to assist medical personnel on early breast cancer detection. The complete EBCD framework consists of the integration of software and hardware (configuration, obtain data sample/s, analogue to digital conversion and scan data saving), preprocessing, detection of breast cancer size and visualization of the output in the 2D and 3D environment. Once the framework is developed, the framework is converted into a standalone executable file (.exe file) in order to make the system flexible and easy to access.

EBCD framework is developed using MSFS method and NB for breast cancer size detection. The fresh data sample will not go through MSFS processes from scratch again (Stage 1 to Stage 4) as the data normalization methods and features extraction methods involved in forming 8-HybridFeature are already selected and identified in this work. Thus, it helps in reducing the time consumption and computational complexity in EBCD framework. The proposed EBCD algorithm is implemented in the UWB system to develop a complete early breast cancer size detection framework. Fig 12 and Fig 13 show the example of visualization layout in the 2D and 3D environment to detect 2 mm tumor at the location of 2.5 mm, 32.5 mm and 50 mm for x, y and z coordinates respectively.

**Fig 12.**
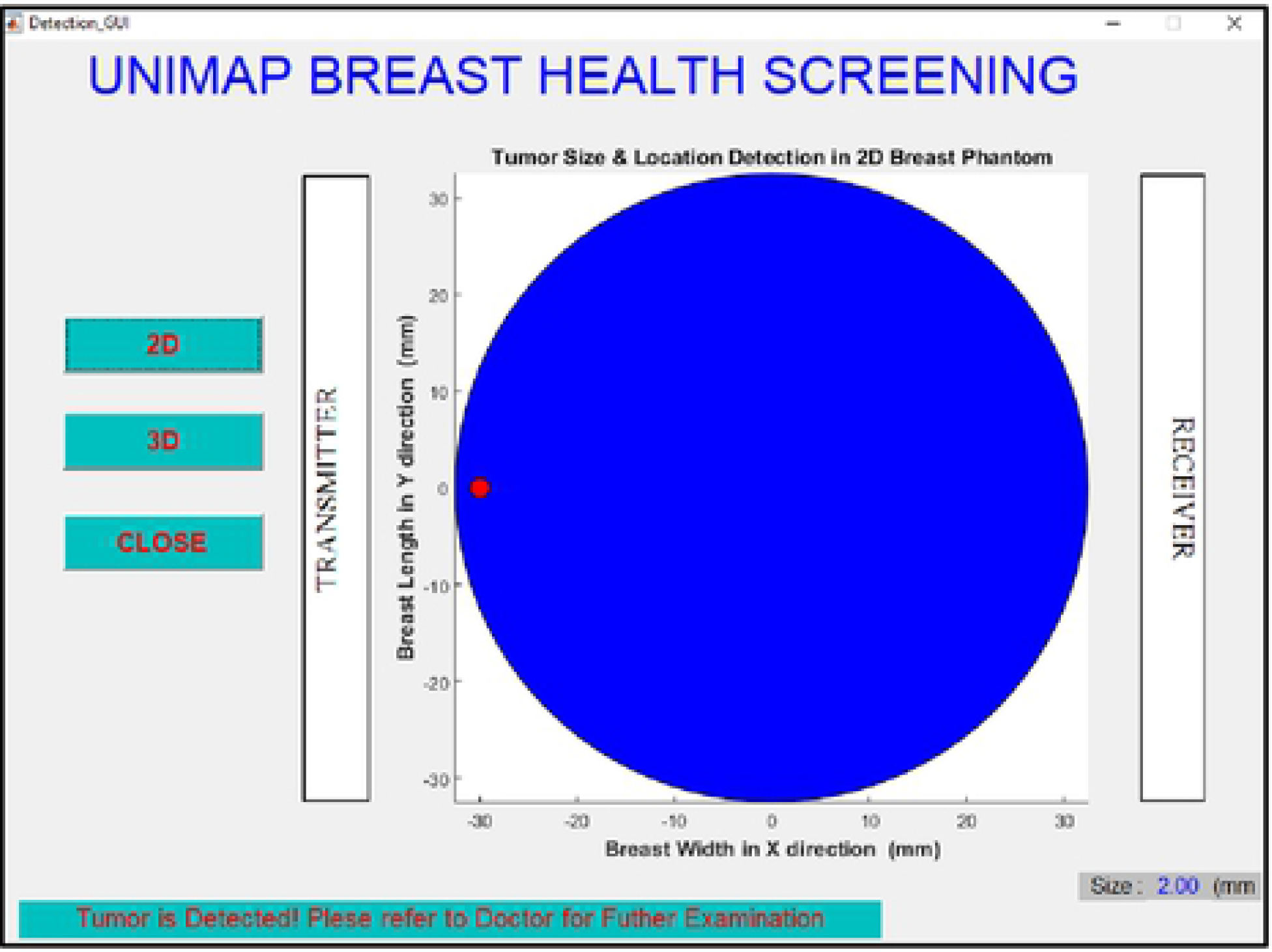
Visualization of Detected 2 mm Tumor Size in 2D Environment

**Fig 13.**
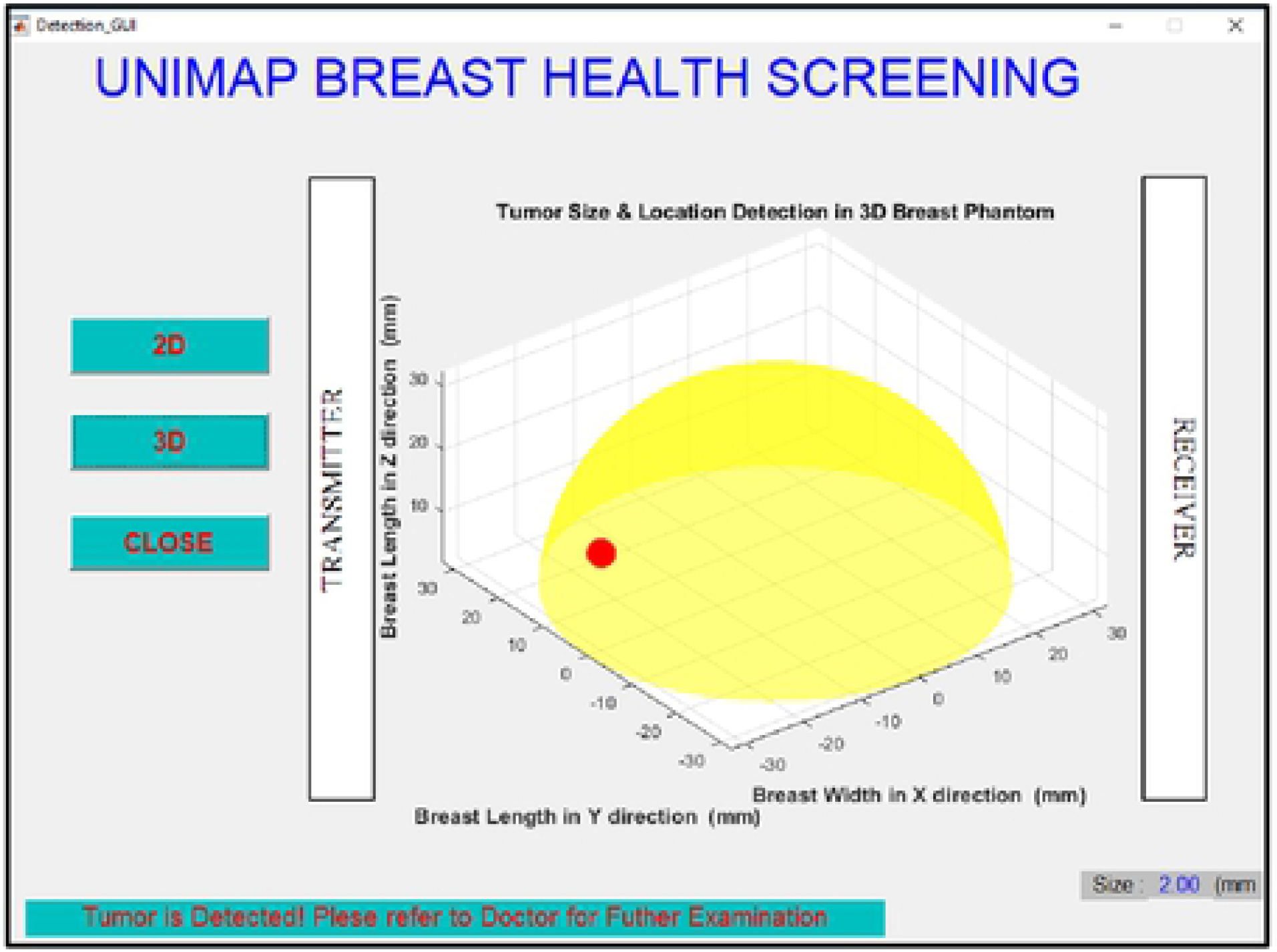
Visualization of Detected 2 mm Tumor Size in 3D Environment

## Conclusions

MSFS method is proposed for early breast cancer detection application. The proposed MSFS method has four stages. The first stage consists of data normalization methods and data reduction. The second stage consist of feature extraction methods, while third stage and fourth stage consist of feature selection and feature fusion, respectively. The selection of data normalization methods and features are done by computing the *p*-value and *F*-value in each stage. The raw data samples go through these stages in order to identify the best data normalization methods, best feature extraction methods and optimum hybrid features. The hybrid features are fused together using feature fusion technique.

Three different datasets (8-HybridFeature, 9-HybridFeature and 10-HybridFeature) are formed through MSFS method. These datasets are tested using three different supervised classifiers (SVM, NB and PNN) to check the robustness of the features. All classifiers obtain classification accuracy of more than 70%. The highest classification accuracy is obtained by 8-HybridFeature dataset tested in NB classifier (91.98%). A complete early breast cancer detection framework is developed. The finalized MSFS methods are implemented in the EBCD framework. The detected size is visualized in the 2D and 3D environment.

## Data Availability Statement

All relevant data are within the paper.

## Acknowledgment

This work is supported by Ministry of Higher Education, Malaysia, Grant FRGS – 9003-00418.

## Author Contributions

Conceived and designed the experiments: VV AMA MJ SK. Performed the experiments: VV SK. Analyzed the data: VV MM TS RAAR. Contributed materials/analysis tools: MJ MNMY RBA. Wrote the paper: VV AMA.

